# The Dendritic Ergic: Microtubule and Actin Cytoskeletons Mediate Stop-and-Go Movement of Mobile Carriers Between Stable Structures

**DOI:** 10.1101/2021.03.31.437880

**Authors:** María de los Ángeles Juricic Urzúa, Javiera Gallardo Rojas, Andrés Couve Correa, Mauricio Cerda, Steffen Härtel Gründler, Carolina González-Silva

**Author notes:** To whom correspondence should be addressed: Carolina González-Silva; Cellular and Molecular Neurobiology Laboratory, Neuroscience Department, Faculty of Medicine, Universidad de Chile, Independencia 1027, Santiago, Chile.

## Abstract

The ER-to-Golgi intermediate compartment (ERGIC) is a membranous organelle that mediates protein transport between the endoplasmic reticulum (ER) and Golgi apparatus. In neurons, clusters of these vesiculotubular structures are situated in throughout the cell in proximity to the ER, passing cargo to the cis-Golgi cisternae located mainly in the perinuclear region. Although ERGIC markers have been identified in neurons, the distribution and dynamics of neuronal ERGIC structures have not been characterized.

Here, we argue that long-distance ERGIC transport occurs via an intermittent mechanism in neurons, with mobile elements moving between stationary structures. Using immunofluorescence microscopy, we detected discrete, irregular ERGIC structures in neural soma and dendrites. Slow live-cell imaging (2 frames/minute; 15 minutes) indicated that 8% of dendritic ERGIC structures were stable, remaining in place over long periods. On the other hand, fast live-cell imaging (2 frames/second; 180 seconds) captured mobile ERGIC structures advancing very short distances along dendrites. Importantly, these distances were consistent with the lengths between the stationary ERGIC structures. Kymography revealed ERGIC elements that moved intermittently, emerging from and fusing with stationary ERGIC structures. Surprisingly, this movement was apparently dependent not only on the integrity of the microtubule cytoskeleton, as has been previously reported, but on the actin cytoskeleton as well.

Our results indicate that the dendritic ERGIC has a dual nature, with both stationary and mobile structures. The neural ERGIC network transports proteins via a stop-and-go movement that is mediated by the microtubule and actin cytoskeletons.

## INTRODUCTION

The ER-to-Golgi intermediate compartment (ERGIC) is a membranous vesiculotubular organelle that extends throughout the cell, functionally mediating bidirectional membrane protein trafficking between the endoplasmic reticulum (ER) and Golgi apparatus (1–6). The lectin protein ERGIC-53 is the chief marker of the ERGIC (7–9).

Two models have been proposed for the role of the ERGIC in ER-to-Golgi transport: the “transport complex” and “stable compartment” models (2). According to the transport complex model, ERGIC structures are transient cargo containers formed by fusion of ER-derived vesicles that migrate and fuse to form cis-Golgi cisternae (2, 10). This model cannot explain why ERGIC structures are preferentially distributed near the ER (1) nor why blocking ERGIC-to-Golgi transport fails to increase the abundance of ERGIC structures (6). On the other hand, the stable compartment model classifies the ERGIC as a separate organelle that receives cargo from the ER and releases anterograde carriers destined for the Golgi (6, 11). In this model, stationary ERGIC structures act as initial post-ER sorting stations. Transport between the ER and Golgi is a two-step process that involves short-range movement from ER exit sites (ERES) to stable ERGIC structures, followed by long-distance transport from the ERGIC to cis-Golgi (2). This model is supported by evidence from live-cell imaging of HeLa cells. Green fluorescent protein (GFP)-ERGIC-53 has been identified in these cells, within stationary vesiculotubular structures that sort ssDsRed (a fluorescent secretory marker protein) into carriers leaving GFP-ERGIC-53-positive structures and moving toward the Golgi (1).

In non-polarized cells, the ERGIC network includes both stationary structures, which remain in a given position over periods of at least 15 minutes, and a population of mobile carriers that allow for transport between the stationary elements (1). Due to the relatively small size and simple morphology of these cells, mobile ERGIC structures only need to cover short distances to connect the ER and Golgi (12).

However, little is known about ERGIC distribution and dynamics in larger cells with more complex morphology, such as neurons (13). Neurons are highly polarized cells with two clearly-differentiated domains: somatodendritic and axonal. Dendrites and axons may extend hundreds of microns away from the soma (13). Secretory organelles must adapt to this complex neuronal morphology; while the ER and functional ERES extend throughout the cell, the Golgi apparatus is restricted to the soma and dendritic Golgi structures. There are at least three specializations of the secretory route in neurons: (i) the Golgi outpost, a discrete Golgi apparatus discontinuous with the somatic Golgi, that is located in the apical dendrites of a subset of neurons (14–18); (ii) the Golgi satellite, consisting of trans-Golgi network structures located in all dendrites (19); and (iii) the spinal apparatus, present in almost all dendritic spines as a membranous structure continuous with dendritic ER, which shares markers, characteristics, and functions with the ER, ERGIC, and Golgi (20–21). It seems that a relatively small proportion of proteins synthesized in the dendritic Golgi are processed locally, leaving the remaining proteins dependent on carriers for retrograde travel. These carriers are presumably ERGIC structures that transport cargo from distant dendritic ERES to somatic Golgi (16).

ERGIC transport structures may be particularly crucial for the local dendritic response to certain environmental stimuli. Such responses may require adjusting the relative abundance of specific plasma membrane proteins, as well as establishment, maintenance, and plastic modifications of synapses.

Although ERGIC markers have been described in neuronal soma and dendrites (15, 20, 22), the distribution, morphology, molecular composition, function, and dynamics of ERGIC structures in these cells remain largely unknown.

Here, we characterize ERGIC distribution and morphology in the somatodendritic domain of hippocampal neurons, using live imaging at two time scales. Additionally, we explore the dynamics of dendritic ERGIC to evaluate whether the transport complex or stable compartment model best explains ERGIC function in neurons.

## RESULTS

### ERGIC distribution, morphology, and relationship to other organelles in the somatodendritic domain

The lectin family protein ERGIC-53 is the most widely-used ERGIC marker (2–3, 6–9, 23–30). Although ERGIC-53 cycles between the ER and Golgi carrying glycosylated proteins (23–24, 27, 31–35), it also accumulates in stationary ERGIC structures (2), and its presence in neurons has been confirmed (20). Therefore, we labelled ERGIC-53 to identify the ERGIC in the somatodendritic domains of cultured rat hippocampal neurons at 14-18 days *in vitro* (DIV). Using immunofluorescence against endogenous ERGIC-53 and confocal microscopy, we detected discrete ERGIC structures spread widely throughout the neuron in the soma as well as the primary, secondary, and tertiary dendritic branches (Fig. 1A).

**Figure 1.**
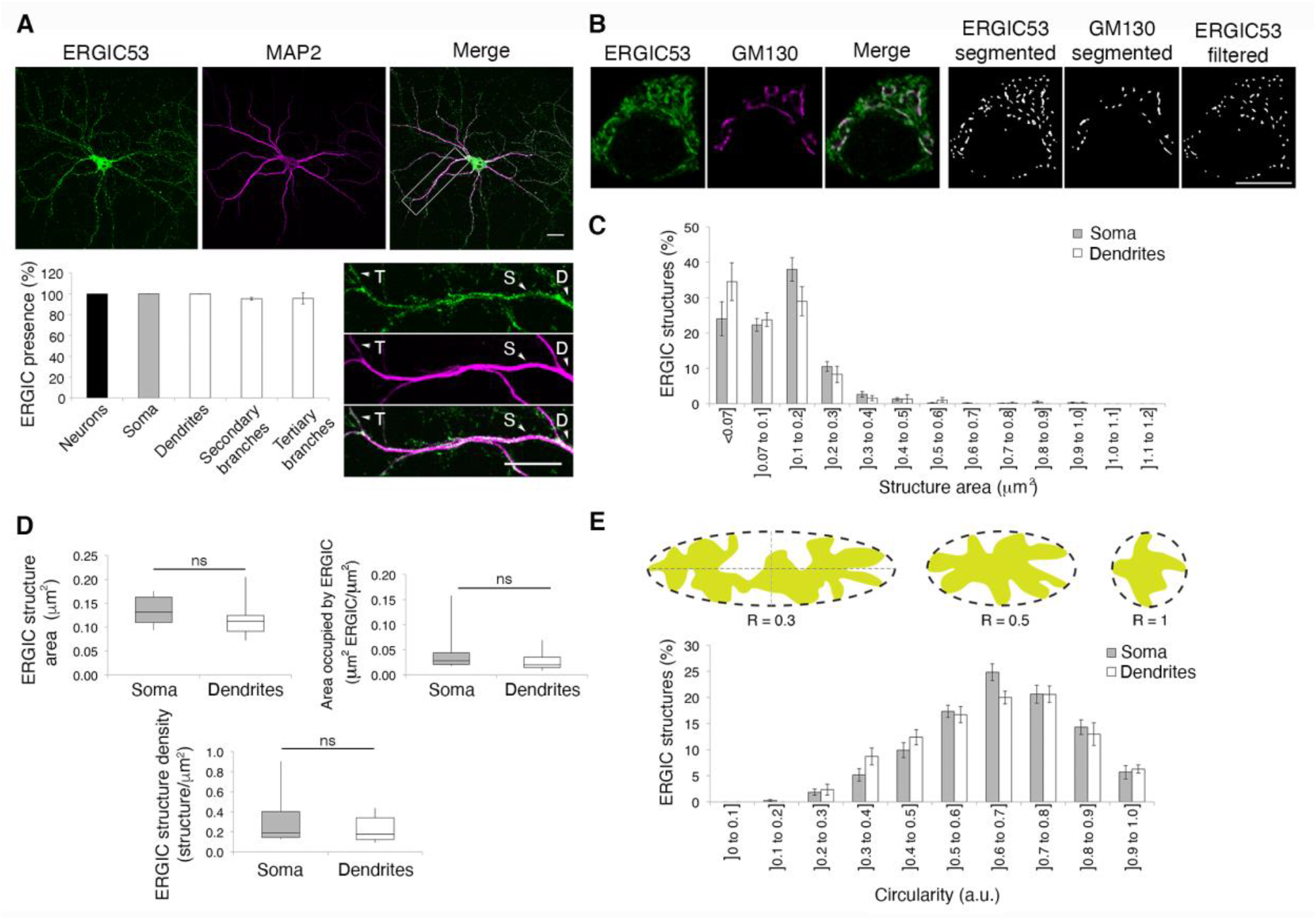
Distribution, size, and morphology of ERGIC structures in the somatodendritic domain. **A) Top.** Immunofluorescence showing endogenous expression of the ERGIC marker ERGIC-53 (green) and somatodendritic domain marker MAP2 (magenta) in cultured rat hippocampal neurons at 14–18 DIV. **Bottom left.** Percentages of neurons, soma, dendrites, and secondary and tertiary branches with ERGIC-53-positive structures. **Bottom right.** Insets from upper row, showing endogenous ERGIC-53 (green) and MAP2 (magenta) expression in dendrites (D) and secondary (S) and tertiary (T) branches. All images were acquired using confocal microscopy (60X objective, no digital zoom); scale bar: 10 μm (3 independent cultures). **B) Left.** Endogenous expression of ERGIC-53 (green) and cis-Golgi marker GM130 (magenta) in soma of cultured rat hippocampal neurons at 14–18 DIV. **Right.** Binary segmented mask of endogenous ERGIC-53 and GM130 expression developed with the local maximum algorithm (see Methods). ERGIC structure area and morphology analyses were performed using this mask. All images were acquired using confocal microscopy (60X objective, 4X digital zoom); scale bar: 10 μm. **C)** Size distribution of ERGIC-53-positive structures in terms of area (μm^2^) in soma and dendrites from cultured rat hippocampal neurons at 14–18 DIV; n=9 neurons (3 independent cultures); scale bar: 10 μm. **D)** Mean ERGIC structure area (μm^2^); relative area occupied by ERGIC structures (μm^2^ ERGIC/μm^2^); and ERGIC structure density (ERGIC structures/μm^2^) in soma and dendrites from neurons in B; n=9 neurons (3 independent cultures). One-sample Wilcoxon signed rank test: n.s.: no significant differences. **E) Top.** ERGIC structure morphology was measured in terms of circularity (see Methods). Examples of various morphologies and corresponding circularities are shown. **Bottom.** ERGIC-53-positive structure morphology distribution in terms of circularity (arbitrary units) in soma and dendrites of neurons in B; n=9 neurons (3 independent cultures).

To characterize the morphology of ERGIC structures, we analyzed confocal microscopy images from immunofluorescence against ERGIC-53 and the cis-Golgi marker GM130. Given the relatively high colocalization of ERGIC-53 and GM130 (Fig. 1B, left panel), we segmented the images and created masks representing ERGIC-53- and GM130-positive structures. We then eliminated all ERGIC-53-positive structures that totally or partially overlapped with GM130, creating a segmented image with the filtered ERGIC-53 structures. This image was used to calculate the size and morphology of this organelle in the somatodendritic domain (Fig. 1B, right panel). A wide distribution of sizes was observed among these ERGIC structures, although most (over 80%) corresponded to structures of less than 0.2 μm^2^ (Fig. 1C). There were no significant differences between the soma and dendrites in terms of the mean area of ERGIC structures, the area occupied by ERGIC structures relative to the total area evaluated, or the density of ERGIC structures (Fig. 1D). Additionally, we analyzed ERGIC morphology in terms of circularity (Fig. 1E, top panel). We found a wide distribution of shapes among ERGIC structures, ranging from tubular (circularity closer to 0) to spherical (circularity closer to 1). Most (over 50%) were vesicular structures with a circularity between 0.5 and 0.8, and a major radius 1.5-2 times greater than the minor radius (Fig. 1E, bottom). There were no significant differences in circularity between the somatic and dendritic ERGIC structures.

To study the relationship of ERGIC structures with other secretory pathway organelles, we analyzed the true colocalization of ERGIC-53 with ER (KDEL) and cis-Golgi (GM130) markers (Fig. 2B and 2C). True colocalization was determined calculating Mander’s coefficients (M1 and M2), subtracting the random component of colocalization (see Methods). Additionally, we evaluated the maximum percentage of true colocalization by determining the true colocalization of ERGIC-53 attached to FITC- (ERGIC-53-FITC) labeled secondary antibody and ERGIC-53 attached to TRITC- (ERGIC-53-TRITC) labeled secondary antibody (Fig. 2A). We obtained true colocalization figures for these markers (M1: 43.7 ± 12.6% in soma and 55.3 ± 1.3% in dendrites; M2: 47.0 ± 11.5% in soma and 40.7 ± 11.3% in dendrites) (Fig. 2D). In the ER, we found almost no overlap in the local distribution of ERGIC-53 and the ER marker KDEL (Fig. 2B and 2D), with a true colocalization below 1% in the soma (M1: 0.3 ± 2.3%; M2: 0.1 ± 2.1%) and below 4% in dendrites (M1: 3.7 ± 4.9%; M2: 3.9 ± 4.9%). Moreover, we found a partial but significant true colocalization of ERGIC-53 and the cis-Golgi marker GM130 (Fig. 2C and 2D). While most ERGIC-53-positive structures were not positive for GM130 (M1: 13.2 ± 9.5% colocalization in soma and 15.1 ± 9.5% in dendrites), a great proportion of GM130-positive structures were also positive for ERGIC-53 (M2: 29.5 ± 17.2% in soma and 32.1 ± 19.0% in dendrites). However, the colocalization of ERGIC-53 and GM130 was significantly lower than that of Rab1A-53-FITC and ERGIC-53-TRITC (p<0.001).

**Figure 2.**
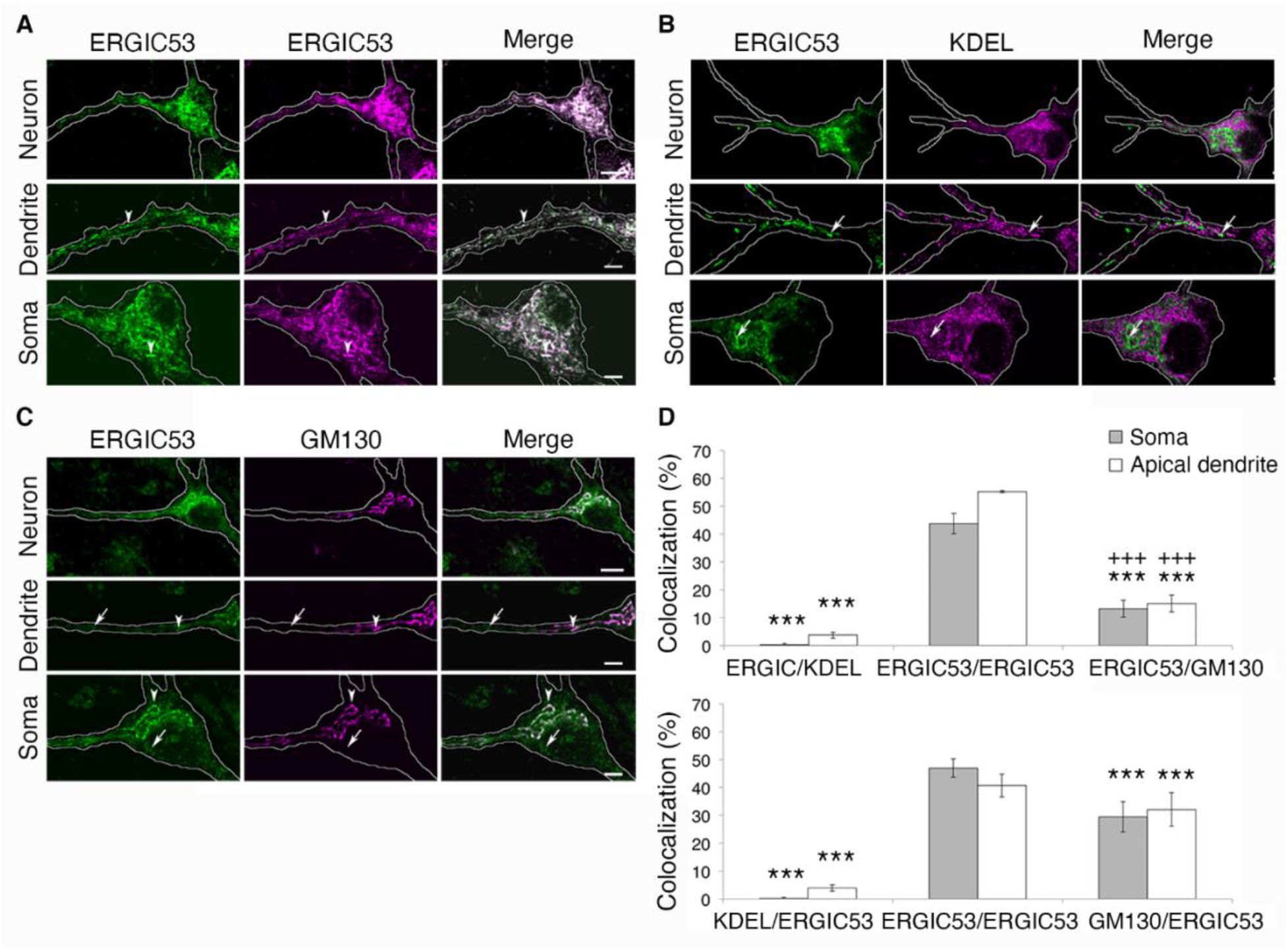
Colocalization of ERGIC structures with ER and Golgi markers in the somatodendritic domain. **A)–C)** Immunofluorescence from cultured rat hippocampal neurons at 14–18 DIV showing endogenous expression of ERGIC-53 attached to FITC-labeled secondary antibody (green) and ERGIC-53 attached to TRITC-labeled secondary antibody (magenta, A), ER marker KDEL (B), and cis-Golgi marker GM130 (C) in neurons, dendrites, and soma. All images were acquired using confocal microscopy (60X objective, 4X digital zoom); scale bars: neuron: 10 μm, dendrite: 5 μm, soma: 5 μm. **D)** True colocalization of ERGIC-53 and ERGIC-53 (n=12 neurons), KDEL (n=17 neurons), or GM130 (n=9 neurons) per Mander’s coefficients M1 (top) and M2 (bottom). Data from 3 independent cultures, shown as median ± standard error. Student’s t-test: *: P<0.05; **P<0.01; ***P<0.001 for true colocalization of ERGIC-53-FITC/ERGIC-53-TRITC; +P<0.05; ++P<0.01; +++P<0.001 for true colocalization percentages per Mander’s coefficients M1 and M2.

We also used Rab1A as an alternative ERGIC marker to study the relationship between neuronal ERGIC and other secretory organelles. Rab1A is a small GTPase that participates in ER-to-Golgi membrane trafficking by tethering COPII vesicles to the ERGIC and cis-Golgi membranes (36). In NRK cells, stationary Rab1A has a distribution very similar to that of ERGIC-53 (37). We transfected cultured rat hippocampal neurons with GFP-Rab1A and then evaluated the cells by immunofluorescence against endogenous ERGIC-53, KDEL, or GM130 (Supp. Fig. 1). We analyzed the percentage of true colocalization of GFP-Rab1A and the other proteins. GFP-Rab1A partially colocalized with ERGIC-53 (M1: 10.9 ± 10.7% in soma and 6.2 ± 5.8% in dendrites; M2: 9.6 ± 10.3% in soma and 7.1 ± 5.7% in dendrites), and these true colocalization percentages were significantly lower than the corresponding percentages for ERGIC-53-FITC and ERGIC-53-TRITC (p<0.001). As with ERGIC-53, the Rab1A-GFP and KDEL distributions showed almost no overlap, with a true colocalization below 4% (M1: 2.2 ± 3.2% in soma and 3.2 ± 5.4% in dendrites; M2: 2.6 ± 2.9% in soma and 3.6 ± 5.1% in dendrites). On the other hand, GFP-Rab1A strongly colocalized with GM130 in the soma (M1: 28.5 ± 8.0%; M2: 60.2 ± 12.5%), at levels comparable to or higher than the colocalization of GM130 and ERGIC-53. As with ERGIC-53, most GM130-positive structures were also GFP-Rab1A-positive, while not all GFP-Rab1A-positive structures were positive for GM130 (Supp. Fig. 1). We were not able to analyze the true colocalization of GFP-Rab1A and GM130 in dendrites because we did not detect any dendritic Golgi in these neurons.

### Two distinct populations of dendritic ERGIC structures: characterization of the mobile ERGIC pool

Live-cell imaging in non-polarized HeLa cells has revealed two distinct types of ERGIC structures: a stationary population that remains in place for at least 15 minutes at a time, and a mobile pool that moves bidirectionally, connecting the stable structures to the ER and Golgi (1). Although this evidence supports the stable compartment model in HeLa cells, it remains unknown whether these findings apply to other cell types, particularly polarized cells such as neurons. Furthermore, we wanted to explore how this stop-and-go movement might enable long-distance transport in dendrites.

Here, we used fluorescence microscopy and live-cell imaging of cultured rat hippocampal neurons at 14-18 DIV transfected with YFP-ERGIC-53 to observe ERGIC dynamics in dendrites. We applied two approaches: “slow” live-cell imaging, recording neurons for 15 minutes at 2 frames/minute (Fig. 3), and “fast” live-cell imaging, recording for 180 seconds at 2 frames/second (Fig. 4).

**Figure 3.**
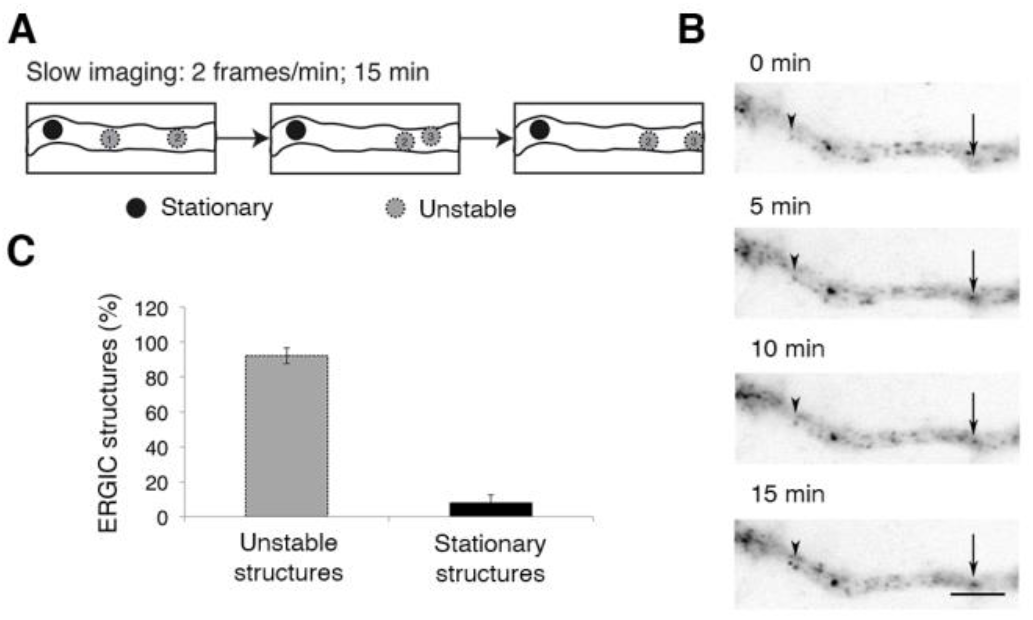
Stationary and unstable dendritic ERGIC structures observed with slow live-cell imaging. Slow live-cell imaging (2 frames/minute) from cultured rat hippocampal neurons at 14–18 DIV transfected with YFP-ERGIC-53. **A)** Scheme showing the types of ERGIC structures identified: stationary structures whose position did not vary significantly during the recording (black circle) and unstable structures that suddenly appeared, disappeared, or changed position significantly (more than 0.512 μm). **B)** Slow live-cell imaging of a dendrite. Stationary (arrowhead) and unstable (arrow) structures were observed. It should be noted that an unstable structure suddenly appeared next to the stationary structure shown at 15 minutes. All images were acquired using fluorescence microscopy with a 100X water-immersion objective lens at a rate of 2 frames per minute for 15 minutes. Scale bar: 5 μm. **C)** Percentage of stationary and unstable ERGIC structures in dendrites. Data: mean ± standard error, n=6 dendrites (6 neurons, 3 independent cultures).

**Figure 4.**
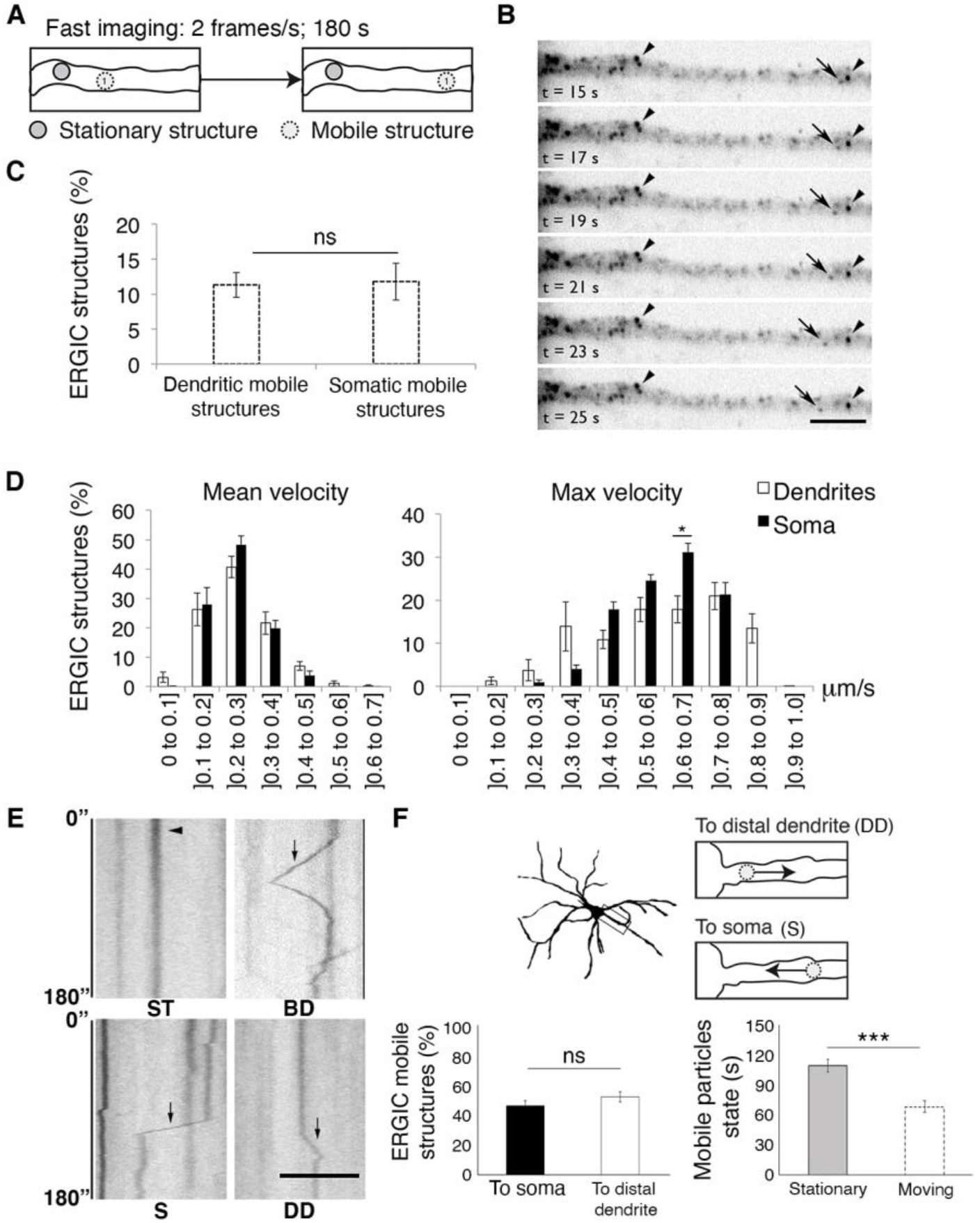
Stationary and mobile dendritic ERGIC structures observed with fast live-cell imaging. Fast live-cell imaging (2 frames/second) from cultured rat hippocampal neurons at 14–18 DIV transfected with YFP-ERGIC-53. **A)** Scheme showing the types of ERGIC structures identified: stationary structures whose position did not vary significantly over time (gray circle with continuous line) and mobile structures that significantly changed position (more than 0.512 μm) during the recording (light gray circle with dotted line). **B)** Fast live-cell imaging of a dendrite. Stationary (arrowhead) and mobile (arrow) structures were observed. All images were acquired using fluorescence microscopy with a 100X water-immersion objective lens at a rate of 2 frames per second for 180 seconds. Scale bar: 5 μm. **C)** Percentage of mobile ERGIC structures in dendrites (n=13, 3 independent cultures) and soma (n=8, 3 independent cultures). Data: mean ± standard error. **D)** Histograms showing mean (μm/second) and maximum velocity (μm/second) of mobile ERGIC structures in soma (black bars) and dendrites (white bars). **E)** Kymographs from mobile ERGIC structures in dendrites (n=13, 3 independent cultures) during fast live-cell imaging records (0 to 180 seconds) showing movement of individual ERGIC structures. The kymographs showed stationary (ST) and mobile structures that moved bidirectionally (BD) or mobile structures with movements directed towards either the soma (S) or distal regions of the dendrites (DD). Scale bar: 5 μm. **F) Top.** Scheme showing mobile ERGIC structures moving towards the soma or distal regions of dendrites. **Bottom left.** Percentage of mobile ERGIC structures in dendrites with movement directed towards the soma or distal dendrites. **Bottom right.** Seconds in which mobile particles were stationary or moving. One-sample Wilcoxon signed rank test: n.s.: no significant differences; *p<0.05; ***p<0.001.

The slow live-cell imaging classified an element as “stationary” when the structure remained in the same position without evident morphological change for the entire 15-minute recording period. An “unstable” element was defined as any structure that disappeared or changed position or shape (Fig. 3A). About 8% of ERGIC structures remained stationary (Fig. 3B and 3C). Most of the unstable structures were untrackable at this recording velocity, likely because they moved too fast to be identified as mobile structures in consecutive frames (see Methods).

We then used “fast” live-cell imaging to observe and characterize mobile ERGIC structures in dendrites. We defined an ERGIC structure as “stationary” if the structure remained in the same position or moved less than 512 nm during the recording (180 seconds, 2 frames/second). A “mobile” ERGIC structure was defined as any structure that changed position by at least in 512 nm (Fig. 4A, see Methods).

We found that about 11% of ERGIC structures in the somatodendritic domain were mobile (11.8 ± 7.5% in soma and 11.3 ± 6.4% in dendrites) (Fig. 4B and 4C). We tracked the movement of mobile ERGIC structures and measured mean and maximum velocity of each structure during the time it was moving (Fig. 4D). We found that over 90% of mobile ERGIC structures had a mean velocity less than or equal to 0.4 μm/s in both the soma and dendrites (Fig. 4D, left). On the other hand, maximum velocity varied widely, ranging from 0.1 to 0.9 μm/s (Fig. 4D, right). Interestingly, dendritic ERGIC structures reached a higher maximum velocity than somatic structures, with dendritic elements moving achieving speeds of 0.9 μm/s (Fig. 4D). Using kymographs from the fast live-cell imaging series, we evaluated the movement of individual ERGIC structures. We observed stationary structures (ST) (Fig. 4E, top left) as well as mobile structures with short- and long-range movements. Movements were directed towards either the soma (S) or distal regions of the dendrites (DD) (Fig. 4E, bottom left and right). In some cases, the same structure moved bidirectionally (BD) (Fig. 4E, top right). No significant preference for any direction was found (48.0 ± 3.9% to soma; 53.1 ± 3.9 to distal dendrites; one-sample Wilcoxon signed rank test: n.s., p>0.5) (Fig. 4F, bottom left). Moreover, mobile particles (directed towards the soma or distal dendrites) spent significantly more time in a stationary state, moving only about 40% of the recorded time (stationary state: 110.76 ± 21.11 s; moving state: 69.24 ± 21.11 s; one-sample Wilcoxon signed rank test: ***p<0.001) (Fig. 4F, bottom right).

### Microtubule and actin cytoskeleton involvement in movement of mobile dendritic ERGIC structures

It has been shown that ERGIC structures move along the microtubule cytoskeleton in non-polarized cells (1). To characterize ERGIC structure movement in neurons, we investigated the role of the microtubule cytoskeleton in dendritic ERGIC transport. We treated cultured rat hippocampal neurons at 14–18 DIV transfected with YFP-ERGIC-53 with nocodazole (12.5 μM, 4 hours) to induce microtubule depolymerization. We studied the movement of ERGIC structures using fast live-cell imaging and observed that the treatment did not affect the percentage of mobile ERGIC structures (Fig. 5A). However, nocodazole significantly increased the number of mobile ERGIC structures moving at lower mean velocities, without affecting maximum velocity (Fig. 5B and 5C). Because dynein has been shown to be involved in the transport of ERGIC structures in non-polarized cells (38), we also studied the role of dynein in dendritic ERGIC transport. We overexpressed dynamitin in cultured rat hippocampal neurons at 14–18 DIV co-transfected with YFP-ERGIC-53. Dynamitin is a dynactin complex protein that allows dynein to interact with the vesicles. Dynamitin overexpression disassembles the dynactin complex, thus interfering with dynein-mediated vesicle transport (38). We studied ERGIC structure movement using fast live-cell imaging and observed that dynamitin overexpression did not affect the percentage of mobile ERGIC structures (Fig. 5D).

**Figure 5.**
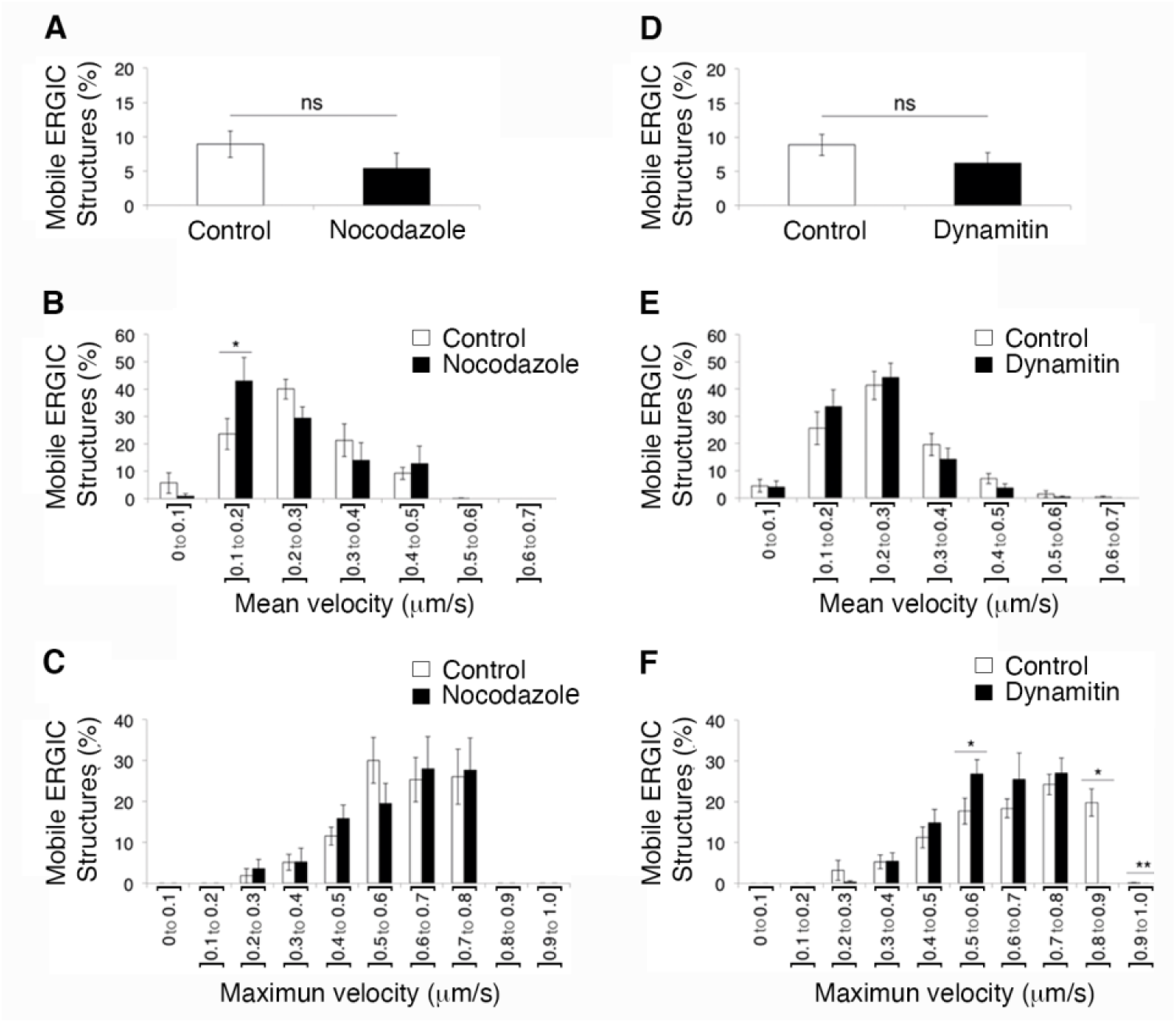
Role of microtubules and dynein in mobile dendritic ERGIC structure transport. Analyses performed using images acquired with fast live-cell imaging (2 frames/second, 180 seconds, 100X objective) of cultured rat hippocampal neurons at 14–18 DIV transfected with YFP-ERGIC-53 and treated with 12.5 μM nocodazole for 4 hours (n=5 neurons) or DMSO as a control (n=8 neurons) (A, B, C), or transfected with YFP-ERGIC-53 and dynamitin-myc (n=5 neurons) or YFP-ERGIC-53 alone as a control (n=4 neurons) (D, E, F). **A)** Percentage of mobile ERGIC structures in neurons treated with DMSO (control) or 12.5 μM nocodazole. **B)** Histograms showing the mean velocity (μm/second) of mobile ERGIC structures in neurons treated with 12.5 μM nocodazole (black bars) or DMSO (control, white bars). **C)** Histograms showing the maximum of velocity (μm/second) of mobile ERGIC structures in neurons treated with 12.5 μM nocodazole (black bars) or DMSO (control, white bars). **D)** Percentage of mobile ERGIC structures in neurons transfected with YFP-ERGIC-53 and dynamitin-myc or YFP-ERGIC-53 alone (control). **E)** Histograms showing the mean velocity (μm/second) of mobile ERGIC structures in neurons transfected with YFP-ERGIC-53 and dynamitin-myc (black bars) or YFP-ERGIC-53 alone (control, white bars). **F)** Histograms showing the maximum velocity (μm/second) of mobile ERGIC structures in neurons transfected with YFP-ERGIC-53 and dynamitin-myc (black bars) or YFP-ERGIC-53 alone (control, white bars). All data presented as mean ± standard error. One-sample Wilcoxon signed rank test: n.s.: no significant differences, *p<0.05, **p<0.01.

In contrast to the nocodazole experiment, however, dynamitin overexpression significantly decreased the maximum velocity of mobile ERGIC structures, without affecting mean velocity (Fig. 5E and 5F). Most notably, no ERGIC structures with velocities over 0.8 μm/second were detected in this condition. These results indicate that the microtubule cytoskeleton and dynein molecular motor are involved in ERGIC structure transport in dendrites. However, these findings do not explain the entire transport process, as some mobile ERGIC structures remained present even at high doses of nocodazole or levels of dynamitin overexpression.

Although no previous studies have described involvement of the actin cytoskeleton in ERGIC structure transport, the finding that some ERGIC structures remained mobile in the presence of high doses nocodazole led us to explore a potential role of this protein. To induce actin cytoskeleton depolymerization, we treated cultured rat hippocampal neurons at 14–18 DIV transfected with YFP-ERGIC-53 with latrunculin (1 μM, 30 minutes). Surprisingly, we observed that latrunculin treatment significantly decreased the percentage of mobile ERGIC structures (from 11.9 ± 4.2% in neurons treated with DMSO to 8.1 ± 3.5% in neurons treated with latrunculin, p<0.005, Fig. 6A) and concomitantly increased the number of ERGIC structures moving at lower mean velocities (p<0.005, Fig. 6B), without affecting maximum velocity (Fig. 6C). These results suggest that that the actin cytoskeleton plays a more significant role in mobile dendritic ERGIC structure transport than the microtubule cytoskeleton.

**Figure 6.**
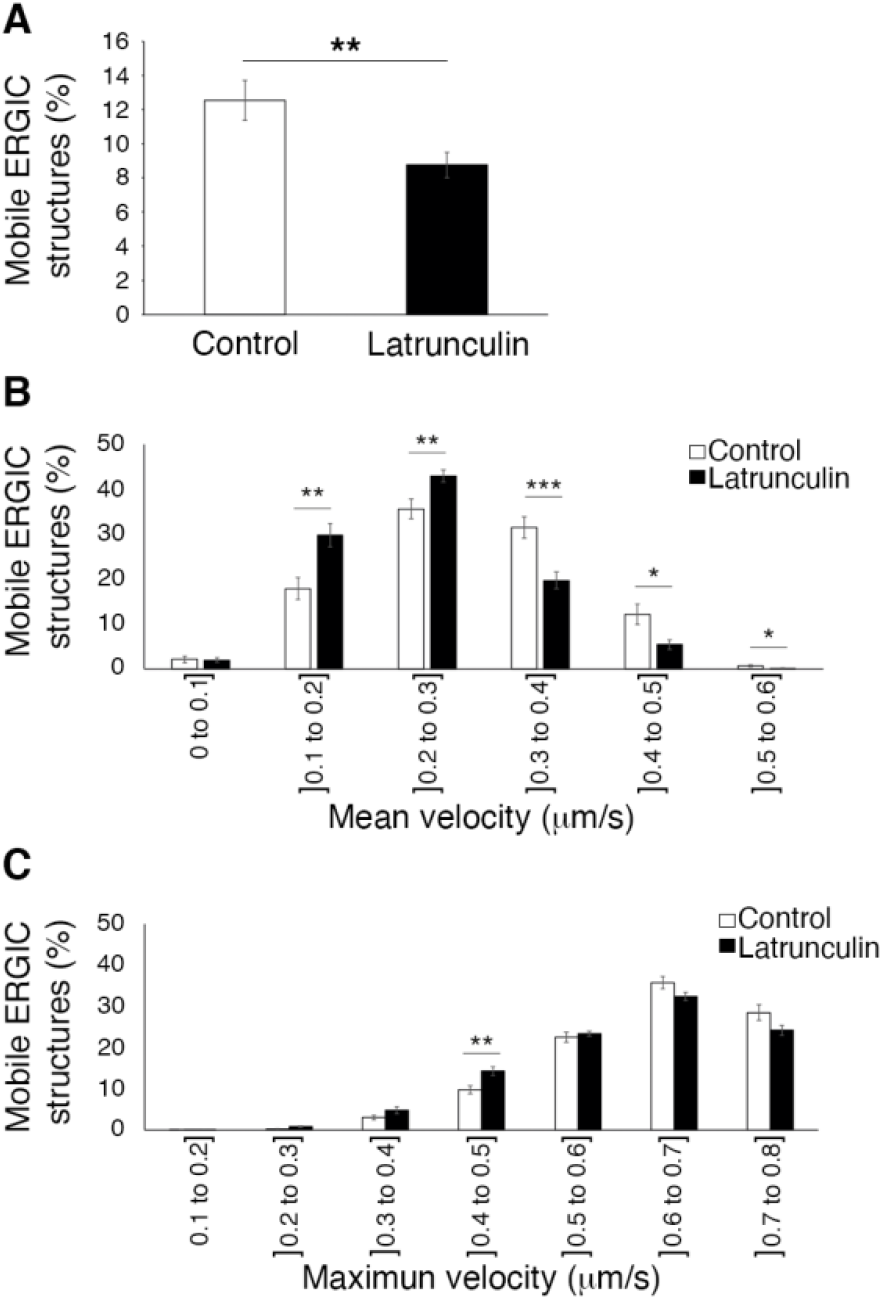
Role of actin in mobile dendritic ERGIC structure transport. Analyses based on fast live-cell imaging (2 frames/second, 180 seconds, 100X objective) of cultured rat hippocampal neurons at 14–18 DIV transfected with YFP-ERGIC-53 and treated with 1 μM latrunculin for 30 minutes (n=22 neurons) or DMSO as a control (n=13 neurons). **A)** Percentage of mobile ERGIC structures in neurons treated with DMSO (control) or 1 μM latrunculin. **B)** Histograms showing the mean velocity (μm/second) of mobile ERGIC structures in neurons treated with latrunculin 1 μM (black bars) or DMSO (control, white bars). **C)** Histograms showing the maximum velocity (μm/second) of mobile ERGIC structures in neurons treated with 1 μM latrunculin (black bars) or DMSO (control, white bars). All data presented as mean ± standard error. One-sample Wilcoxon signed rank test: n.s.: no significant differences, *p<0.05, **p<0.01, ***p<0.005.

### Stop-and-go movement of dendritic ERGIC structures

To characterize the movement of individual mobile ERGIC structures in dendrites as well as their relationship with stable structures, we used kymographs of the fast live-cell imaging series to study mobile ERGIC structures in cultured rat hippocampal neurons at 14–18 DIV transfected with YFP-ERGIC-53 (Fig. 7A). Stop-and-go movement of ERGIC structures and fusion with stationary ERGIC structures were apparent (Fig. 7A). We wondered whether such arrangements could reflect a relay movement of mobile ERGIC structures between nearby static structures (stationary for at least 15 minutes, as defined in the slow live-cell imaging). As it was not possible to simultaneously track stationary and mobile ERGIC structures over long periods, we compared the distances separating adjacent stationary ERGIC structures observed with slow live-cell imaging (Fig. 7B) and the distances traveled by mobile ERGIC structures observed with fast live-cell imaging (Fig. 7C).

**Figure 7.**
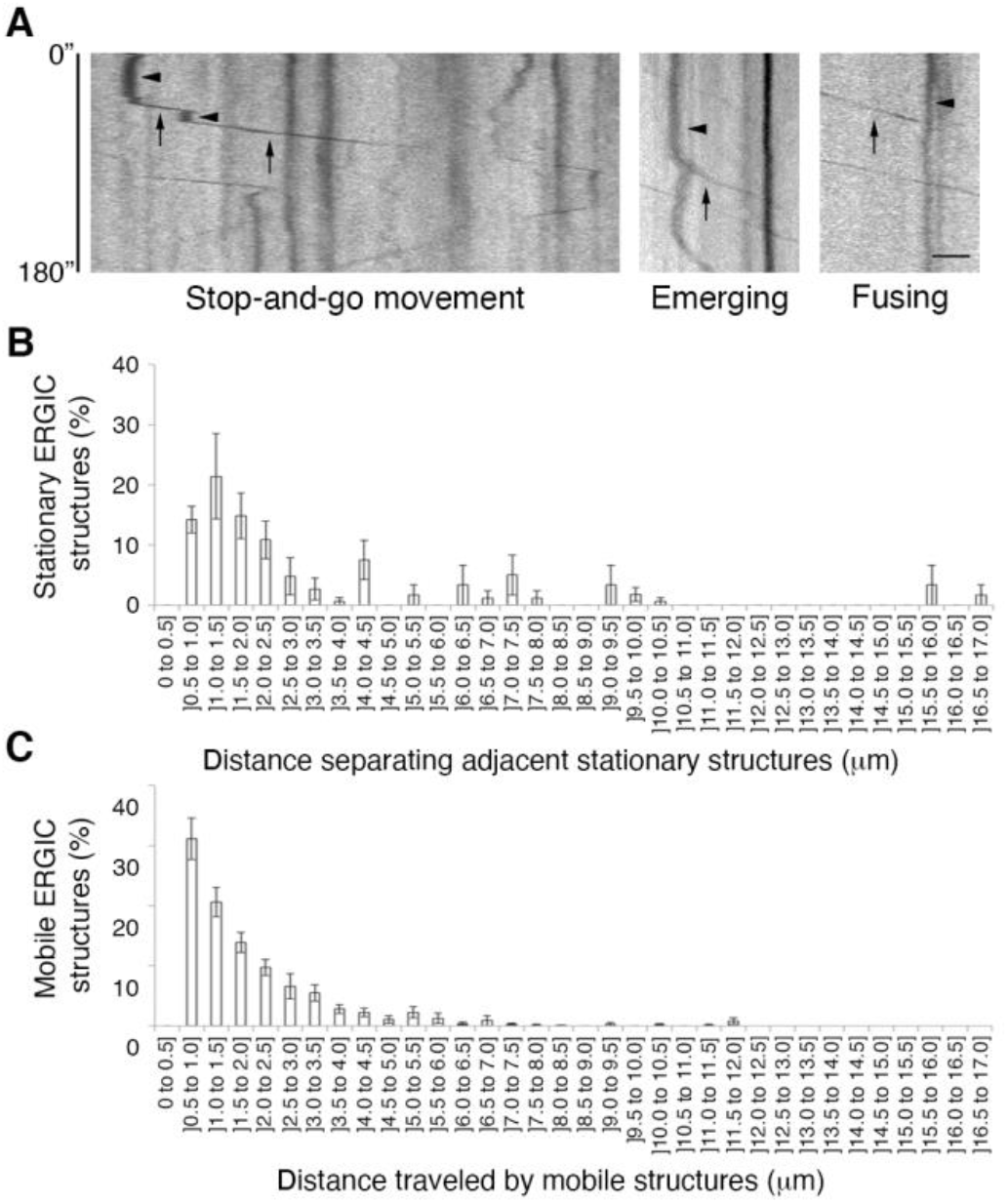
Characterization of ERGIC structure movements in the dendrites. Analyses based on fast live-cell imaging (2 frames/second, 180 seconds, 100X objective) of mobile structures (n=13 dendrites, 3 independent cultures) or slow live-cell imaging (2 frames/minute, 15 minutes, 100X objective) of stationary structures (n=6 dendrites, 3 independent cultures) in cultured rat hippocampal neurons at 14–18 DIV transfected with YFP-ERGIC-53. About 30 kymographs of each dendrite were recorded for each experiment, to analyze the ERGIC across the entire width of the dendrite. **A)** Representative kymographs of mobile dendritic ERGIC structures from fast live-cell imaging (0 to 180 seconds), showing stop-and-go movement of individual ERGIC structures (left), or ERGIC structures emerging from (middle) or fusing with (right) another ERGIC structure. Scale bar: 5 μm. **B)** Histogram showing the distances (μm) separating adjacent stationary ERGIC structures per slow live-cell imaging. **C)** Histogram showing distances (μm) traveled by mobile ERGIC structures per fast live-cell imaging. All data presented as mean ± standard error.

We found that the stationary ERGIC structures were distributed along the entire dendrite. Most stationary structures (82.3%) were separated by distances of less than 5 μm. We found a wide distribution of distances traveled by individual structures, with some structures traveling as far as 12 μm. Interestingly, however, 93.4% of mobile ERGIC structures traveled 5 μm or less (Fig. 7C), supporting the concept of a relay movement.

## DISCUSSION

As in non-neuronal cells (1), neuronal ERGIC is a vesiculotubular organelle that extends throughout the somatodendritic domain, including the secondary and tertiary dendritic branches (Fig. 1).

Dendritic ERGIC structures were defined by the presence of ERGIC-53. Over 80% of the structures identified had a cross section of ~0.2 μm^2^ in confocal images, consistent with previous observations in non-neuronal cells (1).

As our data relied largely on the distribution patterns and dynamics of the most widely-used ERGIC marker, the lectin protein ERGIC-53, we complemented ERGIC-53 colocalization analyses with experiments using alternative organelle markers, such as KDEL (ER) and GM130 (cis-Golgi). As has been previously reported in non-neuronal cells (23), and consistent with the role of ERGIC-53 in transporting proteins to the Golgi apparatus (25), ERGIC-53 localization showed almost no overlap with KDEL localization and partial colocalization with GM130, reflecting its presence in the organelle in its stationary state (Fig. 2). However, other ERGIC markers have also been reported in the literature, including Rab1A (36–37), Rab2 (39), and β-COP (6). To improve our characterization of neuronal ERGIC, we selected Rab1A, as stationary Rab1A has a very similar distribution to ERGIC-53 in NRK cells (37). Surprisingly, we found very different distributions of Rab1A and ERGIC-53 in neurons and low colocalization of the two markers (~10%, Supp. Fig. 1). These results diverge sharply from distribution patterns in other cell types (37) but are quite similar to those observed in PC12 cells. The PC12 cell line is a classical neuronal cell differentiation model in which cells mainly express ERGIC-53 in the soma and Rab1A in neurites, with almost no colocalization between the two markers (40). These expression patterns might reflect differences between neuronal and other cell types. This issue remains to be clarified.

In terms of dynamics, we found that the dendritic ERGIC was composed of both stationary and mobile structures, as in non-polarized cells (1), with stationary structures more predominant than mobile elements. These stationary ERGIC structures, observed by “slow” live cell imaging (2 frames/minute), remained static for at least 15 minutes (Fig. 3). On the other hand, “fast” live cell imaging (2 frames/second) indicated that only ~11% of ERGIC structures were mobile in this time frame (Fig. 4), similar to the 20% figure reported by Ben-Tekaya et al. (2005) under comparable conditions (5 frames/second) in HeLa cells. Furthermore, the dendritic mobile structure velocities (~1 μm/second) were in the range of those observed in non-neuronal cells (~1-7 μm/second). It remains to be determined whether the lower percentage of mobile structures and slower velocities observed are due to different recording rates or represent real differences attributable to the particular characteristics of neurons. Moreover, as in non-neuronal cells (1), mobile dendritic ERGIC structures moved in stop-and-go intervals, emerging from and fusing with stationary ERGIC structures bidirectionally (Fig. 7). Over 90% of mobile dendritic ERGIC structures traveled 5 μm or less, the same distance that separated over 80% of adjacent stationary dendritic ERGIC structures (Fig. 7). These results suggest that mobile ERGIC structures functionally connect adjacent stationary structures in dendrites. Simultaneous live-cell imaging of stationary and mobile dendritic ERGIC structures, using ERGIC markers in conjunction with proteins transported through the ERGIC to other organelles, could be used to evaluate this conjecture.

Taken together, our data supports the “stable compartment” rather than the “transport complex” model, as the most abundant ERGIC structures in the dendrites were stationary over long periods while the mobile structures moved short distances in a stop-and-go fashion, emerging from or fusing with stationary ERGIC structures.

In terms of the role of dendritic ERGIC in connecting the ER to the Golgi, it is important to note that most dendrites have no Golgi outposts (16–17); therefore, most newly-synthesized proteins presumably depend on stable and mobile ERGIC structures for transport from the dendritic ERES to the somatic Golgi apparatus (16). Another possibility is that proteins emerging from the dendritic ERES move through ERGIC structures to the recently-reported Golgi satellite structures (19) or spinal apparatus (20). Horton et al. (2003) showed that over 85% of newly-synthesized proteins leaving the ER from dendritic ERES move in ERGIC-like structures for long-range transport to the somatic Golgi apparatus for further processing, while only 15% are processed locally in dendritic Golgi outposts. The mechanisms underlying the routing of proteins for “somatic” or “local” processing, likely according to the specific characteristics of the protein and/or the needs of the neuron, is a question that remains to be addressed.

The main limitation of our study was fluorophore photostability, which limited the recording rate. Our “fast” live-cell imaging (2 frames/second) was unable to track ERGIC structures for more than 90 seconds, due to photobleaching of YFP-ERGIC-53. While “slow” live-cell imaging (2 frames/minute) allowed us to track stationary structures that remained in the same position, we cannot assume that all of the stationary structures identified in the slow live-cell imaging were the same as those observed in the fast imaging. Further experiments at different time scales would be helpful to complement our results. Moreover, even the “fast” live-cell imaging did not permit us to track structures moving faster than 1.024 μm/second. Therefore, we cannot discard the possibility that there are faster ERGIC structures in neurons that could be affected by nocodazole treatment, as has been previously reported (30, 41–42). It would be helpful to have a live-cell imaging technique that would allow for tracking at a faster recording rate and over longer periods simultaneously, in order to address the full range of dendritic ERGIC structure velocities.

Contrary to previous reports in other cell types (41), only a minor population of mobile dendritic ERGIC structures was sensitive to microtubule cytoskeleton disruption (~15%, see Fig. 5), with only a subpopulation of these elements sensitive to dynein complex destabilization (~10%, see Fig. 5). Moreover, even in these sensitive populations, nocodazole or dynamitin diminished the velocity of dendritic ERGIC structure transport but did not affect the percentage of mobile structures. These results were very surprising and divergent from previous reports in other cell types, where ERGIC transport has been observed to rely mainly on microtubule cytoskeleton integrity (41).

The most exciting contribution of our work is the identification of a group of mobile dendritic ERGIC structures (moving at velocities between 0.3 and 0.6 μm/second) sensitive to actin cytoskeleton depolymerization, as demonstrated by latrunculin treatment (Fig. 6). As far as we know, this actin-dependent transport of ERGIC structures has not been previously reported in any cell type. This transport may be related to the well-documented role of the actin cytoskeleton in dendritic spine transport (43) or to the fact that myosin particles have been localized in organelles previously related to movement along microtubules (46). Myosin’s I, VI, V, and II have been associated with endosomes and lysosomes, the Golgi apparatus, ER, and vesicles involved in transport between Golgi compartments, respectively (47–50). Interestingly, Kuznetsov et al. (51) previously reported that organelles in the squid giant axon switch between actin filaments and microtubules during fast axonal transport. The above findings suggest that dendritic ERGIC also moves through both the actin and microtubule cytoskeletons, as we describe here.

In conclusion, our results verify the presence of ERGIC structures throughout the somatodendritic domain and provide evidence of stationary and mobile dendritic ERGIC structures that move in a stop-and-go fashion. This transport is dependent not only on the microtubule cytoskeleton, but, surprisingly, on the actin cytoskeleton as well. Additional studies are need to clarify the role of both cytoskeletons in this transport and to determine the role of stable dendritic ERGIC structures in protein transport, either as an intra-ERGIC or ERGIC-Golgi relay station or as an organelle with a direct secretory function.

## METHODS

### Animals, hippocampal neuron cultures, and transfection

Adult pregnant female Sprague-Dawley rats were purchased from the Central Animal Facility at Universidad Católica de Chile or the Central Animal Facility at Facultad de Medicina, Universidad de Chile. Animals were euthanized by cervical dislocation according to the Guide for Care and Use of Laboratory Animals (The National Academy of Sciences, 1996) and E18 embryos were used for hippocampal neuron cultures. The experimental protocol was approved by the Institutional Bioethics Committee (Facultad de Medicina, Universidad de Chile, CBA # 0655 FMUCH).

Primary hippocampal neurons were cultured from E18 rats according to established procedures (44). Neurons were maintained in Neurobasal Medium (NB) supplemented with 2% B27, glutamine 1 mM, D-Glucose 0.6% v/v, penicillin-streptomycin (10000 units/ml), and amphotericin B 2.5 μg/mL in a 5% CO_2_ humidified incubator at 37°C. These reagents were from GIBCO (Thermo Fisher Scientific, Waltham, MA, USA), Thermo Fisher Scientific (Rockford, IL, USA), and Sigma-Aldrich (St. Louis, MO, USA). Neurons at 10–18 days *in vitro* (DIV) were used for all experiments.

When necessary, neurons (10–18 DIV) were transfected with DNA plasmids as described below using Lipofectamine 2000 Transfection Reagent (Invitrogen, Carlsbad, CA, USA) according to manufacturer instructions. pmRFP-C1 plasmid was obtained from BD Clontech (Palo Alto, CA, USA). YFP-ERGIC-53/p58 plasmid was kindly provided by J. Lippincott-Schwartz (Janelia Research Campus, Ashburn, VA, USA), described in Ward et al. (2001) (40). GFP-Rab1A plasmid was kindly provided by J. Saraste (University of Bergen, Bergen, Norway), described in Sannerud et al. (2006) (41). p50-HA plasmid was kindly provided by T. Schroer (Johns Hopkins University, Baltimore, MD, USA). Experiments were conducted 24 hours after transfection.

### Immunofluorescence

Immunofluorescence was performed under permeabilized conditions as previously described (45). Primary antibodies were the following: rabbit polyclonal anti-ERGIC-53/p58 and mouse anti-MAP2A purchased from Sigma-Aldrich (St. Louis, MO, USA) (code: E1031 and AB5622, respectively), mouse monoclonal anti-KDEL from Enzo Life Sciences (Biocant, Estación Central, SCL, Chile) (code: ADI-SPA-827), rabbit polyclonal anti-KDEL from Thermo Fisher Scientific (Rockford, IL, USA) (code: PAI-013), mouse monoclonal anti-GM130 from BD Biosciences (Franklin Lakes, NJ, USA) (code: 610822), and rabbit polyclonal anti-TGN38 from Bio-Rad (Hercules, CA, USA) (code: AHP1597). Secondary polyclonal antibodies were purchased from Jackson ImmunoResearch Laboratories (West Grove, PA, USA): FITC anti-Rabbit IgG, TRITC anti-Rabbit IgG, Cy5 anti-Rabbit IgG, FITC anti-Mouse IgG, TRITC anti-Mouse IgG, and Cy% anti-Mouse IgG (codes: 711-095-152, 711-025-152, 711-175-152, 715-095-150, 715-025-150, and 715-175-150, respectively).

### Stained-cell imaging, colocalization, and morphological analysis

Images were obtained using an Olympus FluoView FV 1000 confocal laser scanning microscope and a Universal Plan Super Fluor 60X (oil immersion) apochromatic objective lens (Olympus, Center Valley, PA, USA). A 4X digital zoom was used for colocalization and morphologic analysis of ERGIC structures. Raw images were deconvolved using Huygens Scripting (Scientific Volume Imaging, Hilversum, Holland).

A representative central plane from a soma, dendrite, and dendritic branch was selected for each neuron for the experiments characterizing ERGIC distribution in the somatodendritic domain. The percentages of neurons, soma, dendrites, and secondary and tertiary branches containing ERGIC-53/p58-positive structures were manually counted by an independent observer.

For the colocalization analysis, images from seven consecutives planes on the z-axis separated by 200 nm were acquired for each neuronal domain (soma or dendrite). The true colocalization analysis was performed as previously described (52) using the confined displacement algorithm. ImageJ 2.0.0-rc-43/1.50e (NIH, Bethesda, MD, USA) was used for segmentation and IDL Workbench 7.0 (ITT Visual Information Solutions, Eclipse Foundation Inc., Boulder, CO, USA) to calculate true colocalization percentages.

For the morphological analysis of ERGIC structures, a local maxima algorithm was developed using MATLAB R2012a (MathWorks, Natick, MD, USA). Briefly, a normalized maximum-intensity projection was developed based on the seven ERGIC 8-bit Z-stack slides, segmenting by identifying local intensity maxima (MATLAB imhmax function with a parameter of 13%). Segmentations were cleaned by keeping objects with a maximum intensity of at least 15% and a 2-pixel distance from GM130 regions, applying a region-growing algorithm (maximum distance 20 pixels) to account for realistic sizes and avoid merging objects. All parameters remained fixed during image analysis. The segmented images were then analyzed using the ImageJ 2.0.0-rc-43/1.50e *analyze particles* tool (NIH, Bethesda, MD, USA) to calculate parameters such as number of structures, area (μm^2^), and the major and minor axes of the fitted ellipse. These data were used to calculate the area occupied by ERGIC structures (μm^2^ ERGIC/μm^2^) and ERGIC structure density (number of ERGIC structures/μm^2^) in the soma and dendrites. Circularity was calculated as defined in ImageJ (4*area/π*major_axis^2^).

### Live-cell imaging, particle tracking, and kymographs

For live-cell imaging experiments, neurons at 10–18 DIV were transfected with YFP-ERGIC-53/p58 and/or pmRFP-C1 and/or p50-HA. Neurons were placed in a chamber filled with Tyrode’s medium (NaCl 124 mM, KCl 5 mM, CaCl2 2 mM, MgCl2 1mM, D-glucose 30 mM, HEPES 25 mM). Images were obtained using an Olympus BX61W1 spinning microscope and a LUMPlanF1 100X water-immersion objective lens (Olympus, Center Valley, PA, USA). Sequential images were acquired using cell^R (Olympus, Tokyo, Japan) at a temporal resolution of 2 frames/minute for 15 minutes for “slow” live-cell imaging or 2 frames/second for 180 seconds for “fast” live-cell imaging.

To track ERGIC structures using “slow” or “fast” live-cell imaging, a customized particle tracking algorithm was implemented in MATLAB v9.5.0 (The MathWorks, Inc.) based on previously-reported methods (53). The base algorithm minimizes global displacement and provides an optimal strategy for Brownian motion. The tracking algorithm used 3 main parameters: minimal object diameter (4 pixels; 0.256 μm), minimal object intensity (4 arbitrary units), maximal particle instantaneous speed (8 pixels/frame; 0.512 μm/s). Our customization split trajectories into slow/fast using a fixed threshold of 8 pixels (0.512 μm) for particle displacement in the full movie.

Kymographs were used as a graphic representation of the spatial position of dendritic ERGIC structures over time along a longitudinal line drawn parallel to the upper edge of the dendrite. Kymographs from “slow” and “fast” live-cell imaging of dendrites were constructed using the ImageJ 2.0.0-rc-43/1.50e Multi Kymograph tool (NIH, Bethesda, MD, USA). For each dendrite, 30 to 40 kymographs were constructed, moving the original line up or down in 1-pixel intervals to cover the entire width of the dendrite. These kymographs were used to calculate the distance between two adjacent stationary structures (μm), the distance traveled by a mobile structure (μm), and the direction of movement of a mobile structure (towards the soma or distal dendrites).

For the nocodazole treatment experiments, cultured rat hippocampal neurons at 14–18 DIV transfected with YFP-ERGIC-53 were incubated with 12.5 μM nocodazole or DMSO 0.016% v/v (as a control) for 4 hours prior to the experiment. Nocodazole was purchased from Sigma-Aldrich (St. Louis, MO, USA). For the latrunculin treatment, cultured rat hippocampal neurons at 14–18 DIV transfected with YFP-ERGIC-53 were incubated with 1 μM latrunculin or DMSO 0.016% v/v (as a control) for 30 minutes prior to the experiment. Latrunculin A (ab144290) was purchased from Abcam (Cambridge, UK).

### Statistical analysis

All data are presented as mean ± standard error (S.E.). Student’s t-tests or Wilcoxon-Mann-Whitney tests were used to compare means or medians depending on the distribution of the data.

## Data availability

Data will be shared upon request by email to Dr. María de los Ángeles Juricic.

## Author contributions

M.J. designed and performed most of the experiments, interpreted and analyzed data, and wrote the article manuscript. J.G. performed some experiments, analyzed data, and contributed to the drafting of the manuscript. A.C. designed the experiments and interpreted the analyzed data. M.C. and S.H. designed the image analysis algorithm for ERGIC structure morphology and dynamics. C.G. contributed to the drafting, review, and editing of the manuscript. All authors reviewed and approved the final version of this article

## SUPPORTING INFORMATION

**Supp. Fig. 1.**
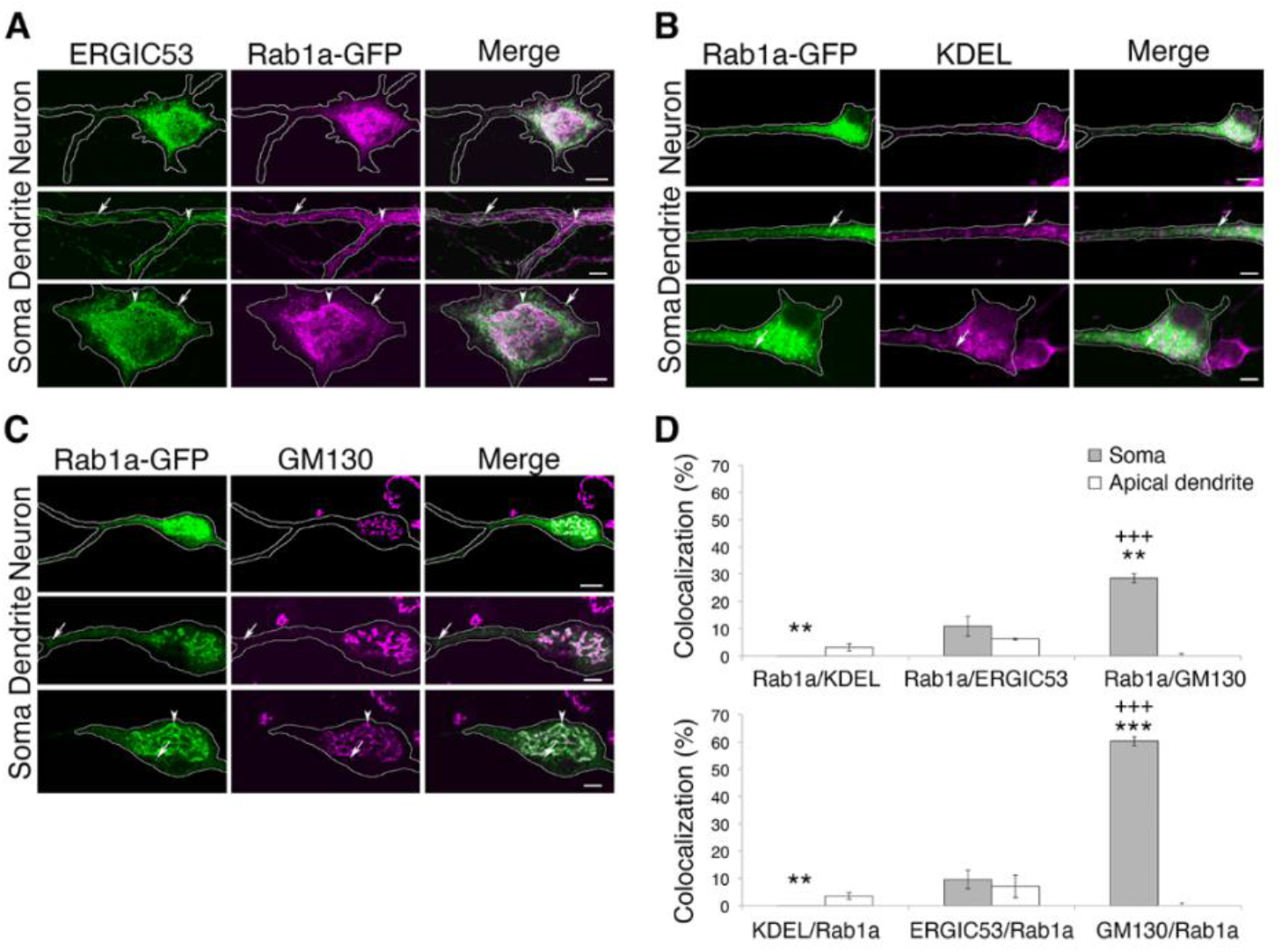
Colocalization of ERGIC structures with Rab1a, KDEL, and GM130 in the somatodendritic domain. **A) to C)** Immunofluorescence from cultured rat hippocampal neurons at 14-18 DIV transfected with GFP-Rab1A (green) showing endogenous expression of ERGIC-53 (A), KDEL (B), and GM130 (C) in neurons (top), dendrites, and soma. All images were acquired by confocal microscopy (60X objective, 4X digital zoom); scale bars: neuron: 10 μm, dendrite: 5 μm, soma: 5 μm. **D)** True colocalization percentages of GFP-Rab1A and ERGIC-53 (n=9 neurons), KDEL (n=7 neurons), or GM130 (n=8 neurons) per Mander’s coefficients M1 (top) and M2 (bottom). Student’s t-test: *: comparisons with true colocalization of ERGIC-53-FITC and GFP-Rab1A (D) +: comparisons between true colocalization percentages per Mander’s coefficients M1 and M2; *p<0.05; **p<0.01; ***p<0.001; +p<0.05; ++p<0.01; +++p<0.001.

